# Oncogenic truncations of ASXL1 enhance a motif for BRD4 ET-domain binding

**DOI:** 10.1101/2021.07.15.452438

**Authors:** Abigail E. Burgess, Torsten Kleffmann, Peter D. Mace

## Abstract

Proper regulation of gene-expression relies on specific protein-protein interactions between a myriad of epigenetic regulators. As such, mutation of genes encoding epigenetic regulators often drive cancer and developmental disorders. Additional sex combs-like protein 1 (ASXL1) is a key example, where mutations frequently drive haematological cancers and can cause developmental disorders. It has been reported that nonsense mutations in ASXL1 promote an interaction with BRD4, another central epigenetic regulator. Here we provide a molecular mechanism for the BRD4-ASXL1 interaction, demonstrating that a motif near to common truncation breakpoints of ASXL1 contains an epitope that binds the ET domain within BRD4. Binding-studies show that this interaction is analogous to common ET-binding modes of BRD4-interactors, and that all three ASX-like protein orthologs (ASXL1–3) contain a functional ET-domain-binding epitope. Crucially, we observe that BRD4-ASXL1 binding is markedly increased in the prevalent ASXL1^Y591X^ truncation that maintains the BRD4-binding epitope, relative to full-length ASXL1 or truncated proteins that delete the epitope. Together, these results show that ASXL1 truncation enhances BRD4 recruitment to transcriptional complexes via its ET domain, which could misdirect regulatory activity of either BRD4 or ASXL1 and may inform potential therapeutic interventions.

## INTRODUCTION

Epigenetic regulation of gene expression has the ability to effect widespread transcriptional changes. Histone modification is a particularly impactful form of epigenetic modification, whereby the addition or removal of posttranslational modifications on histones proteins can change accessibility of chromatin to transcription machinery, or recruit proteins to activate or repress gene expression. Consequently, mutations in genes encoding histone-modifying proteins or proteins that regulate the assembly of histone modification complexes frequently cause cancer, developmental disorders, or other diseases driven by disrupted gene transcription.

One of the most prevalent forms of histone modification is ubiquitination of the C-terminal tail of histone protein H2A (H2AK119Ub, in humans). This histone mark is added by the polycomb repressive complex 1 (PRC1), a ubiquitin-ligase complex that can have variable composition depending on cellular circumstance (Reddington *et al*, 2020; Laugesen *et al*, 2019; Chittock *et al*, 2017). The removal of H2AK119Ub is of equal importance, and is executed by the polycomb repressive deubiquitinase (PR-DUB). The PR-DUB in humans is composed of a catalytic deubiquitinase protein (BAP1 in humans; Calypso in Drosophila), and an Asx-like (ASXL) protein that activates the complex (Scheuermann *et al*, 2010; Sahtoe *et al*, 2016; Foglizzo *et al*, 2018; De *et al*, 2018). There is a single protein, Asx in Drosophila that activates the PR-DUB, but three different ASX-like proteins (ASXL1–3) in humans that can play similar roles (Scheuermann *et al*, 2010). Mutations in either BAP1, or ASXL1 occur at high-frequencies in cancers. Whereas BAP1 mutations drive BAP1 tumour predisposition syndrome, mesotheliomas, melanomas and other neoplasms (Carbone *et al*, 2013; Wiesner *et al*, 2011; Testa *et al*, 2011; Bott *et al*, 2011); ASXL1 mutations are particularly prevalent in myeloid cancers, and are associated with poor prognosis (Balasubramani *et al*, 2015; Gelsi-Boyer *et al*, 2012; Abdel-Wahab *et al*, 2012; Micol *et al*, 2017; Asada *et al*, 2018). Despite their clinical relevance, the exact role of different oncogenic PR-DUB mutations is unclear, and it remains to be determined how prognostic value can be leveraged into targeted therapies for patients with specific mutations.

The domain structure of ASX-like proteins consists of an N-terminal HARE-HTH domain, a central Deubad domain that activates BAP1, and a C-terminal PHD domain posited to bind histone marks. There is a considerable span between the Deubad and C-terminal PHD domain, where the majority of pathological ASXL1 mutations occur (Figure 1A). ASXL1 mutations are mainly comprised of nonsense or frameshift mutations occurring in the 12^th^ exon of ASXL1 and result in a truncation of the protein, terminating before the C-terminal portion of the protein that includes the PHD domain. Such mutations are significantly enriched in acute myeloid leukaemias and myelodysplastic syndromes (Inoue *et al*, 2013; Gelsi-Boyer *et al*, 2012; Asada *et al*, 2019; Gao *et al*, 2013). In addition, inherited truncation variants of a similar length cause a developmental condition known as Bohring-Opitz Syndrome, which is characterised by severe developmental delays, failure to thrive, microcephaly and a set of characteristic facial features (Hoischen *et al*, 2011). Truncations of ASXL3 has also been identified in patients affected by Bainbridge-Ropers Syndrome (Bainbridge et al, 2013), and ASXL2 mutations lead to a related clinically distinct condition (Shashi *et al*, 2016).

**Figure 1.**
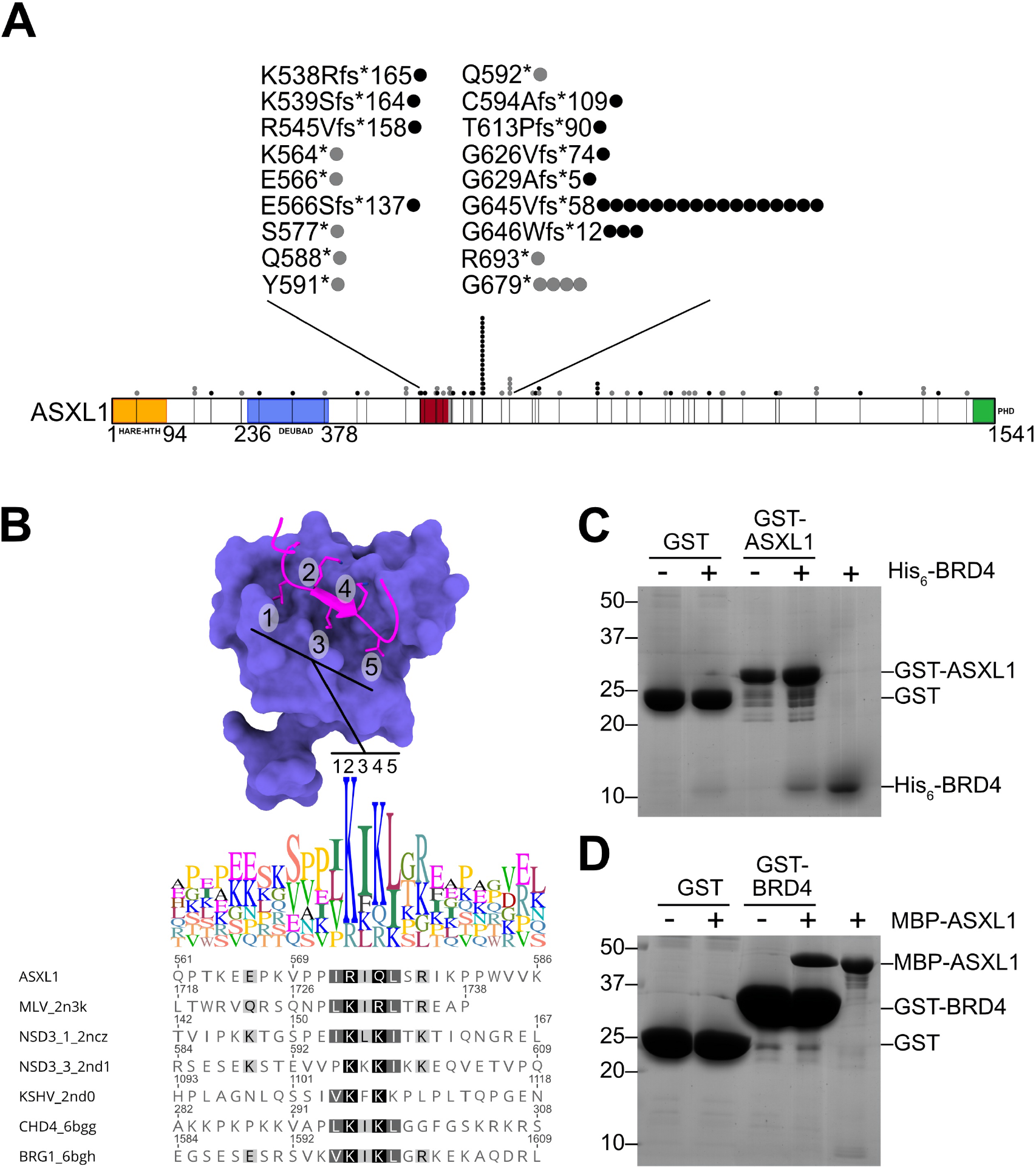
ASXL1 binds the BRD4-ET domain. (A) Schematic representing domain structure of ASXL1. Indicated on ASXL1 is the proposed BRD4 binding site (538-578; dark red) and the relative positions of commonly mutated residues in cancer that produce a truncated protein either by frameshift (black circle) or nonsense (grey circle). (B) Multiple sequence alignment of the BRD4 binding motif in ASXL1 compared with known BRD4-ET interactors, highlighting a conserved five residue motif. These residues we propose bind BRD4 in the same groove of the BRD4-ET as known interactors (PDB: 6BGG; ET bound by CHD4) (C/D) Coomassie-stained gels showing pulldown of His_6_BRD4^600-678^ by GST-ASXL1^537-587^ (C), and the reciprocal pulldown; His_6_MBP-ASXL1^537-587^ by GST-BRD4^600-678^ (D).

Various studies have shown that truncating mutations in ASXL1 lead to genome-wide changes in H2AK119Ub and chromatin status (Asada *et al*, 2018; Balasubramani *et al*, 2015; Nagase *et al*, 2018; Abdel-Wahab *et al*, 2012). However, there has been some debate as to whether truncation of ASXL1 cause a gain- or loss-of-function. Some work indicates that truncated gene-products are unstable proteins and poorly expressed, suggesting loss-of-function (Abdel-Wahab *et al*, 2012), while others have observed a gain-of-function leading to global erasure of H2AK119Ub (Inoue *et al*, 2016; Balasubramani *et al*, 2015). Mechanistically, truncation of ASXL1 appears to affect the ubiquitination status of ASXL1 itself, and subsequently activity of PR-DUB (Daou *et al*, 2018; Asada *et al*, 2018). Recently, a study of a haemopoietic ASXL1^Y588X^ mouse model captured an additional characteristic of ASXL1 driven myeloid malignancies — highlighting a potential gain-of-function interaction between truncated ASXL1 and BRD4, another epigenetic regulator (Yang *et al*, 2018). BRD4 is one of three members of the bromodomain and extra-terminal domain containing (BET) protein family. BRD2, BRD3 and BRD4 are each characterised by two bromodomains nearer their N-terminus, which mediate interaction with acetylated lysine residues, followed by an extra-terminal (ET) domain. BET protein BRD4 in particular is known to be important for chromatin stability and cell cycle progression (Devaiah *et al*, 2016; Dey *et al*, 2009; Mochizuki *et al*, 2008; Yang *et al*, 2008). Because of their role in binding acetylated histone tails and scaffolding regulatory complexes, small-molecule inhibitors that antagonise BET bromodomains have been actively pursued as a means to therapeutically modulate epigenetic gene regulation (Duan *et al*, 2018). The availability of such inhibitors allowed Yang *et al* to show that BET inhibitors can alleviate dysregulation of specific genes downstream of ASXL1 truncation (Yang *et al*, 2018).

Although a gain-of-function interaction with BRD4 is a promising lead in therapeutically targeting ASXL-truncated cancers specifically, the exact mechanism has remained unclear. Here we present biochemical characterisation of an epitope present in all three ASXL proteins that binds ET-domains from BRD2–3. The ASXL1 epitope appears to bind the ET domain tightly compared to other characterised ET-binding motifs. While this work was in preparation, a recent report suggested that ASXL3, but not ASXL1, contains an ET-domain binding motif, which is responsible for linking BRD4 to BAP1 (Szczepanski *et al*, 2020). Our cellular studies using biotin labelling and mass-spectrometry suggest that the ASXL1 ET-binding motif is highly functional in a common oncogenic truncation variant of ASXL1, and is functional but less effective in the context of full-length ASXL1. This work suggests a clear mechanism for gain-of-function of ASXL truncating mutants, and the potential for differential crosstalk between the PR-DUB complexes and BET proteins in either pathological contexts or different cellular contexts.

## RESULTS

### Nonsense frameshift mutations flank an ET-domain binding-motif in ASXL1

ASXL1 (additional sex-comb like-1) truncating mutations occur at high-frequencies in myeloid cancers and are associated with poor prognosis (Gelsi-Boyer *et al*, 2012). The recent finding that ASXL1^aa1-587^ induces a gain-of-function with respect to recruitment of BET bromodomain-containing protein 4 (BRD4) prompted us to investigate the molecular mechanism for the interaction. Performing multiple sequence-alignments we discovered a conserved motif within residues 572–576 of ASXL1 that resembled the sequence used by multiple other proteins to bind to the ET domain of BRD family proteins (Figure 1B). While truncating mutations occur throughout the ASXL1 gene (Figure 1A), there is a striking enrichment of truncating frameshift variants that occur C-terminal to the potential motif at residues 571–576, most prominently at the 645*, 646*12 and 679* variants. The combination of sequence similarity, and proximity to breakpoints of common variants prompted us to test if this region of ASXL1 is sufficient to elicit ET-domain binding.

We first tested the ability of GST-fused ASXL1^537-587^ to bind recombinant BRD4-ET domain. There was a clear interaction between the two proteins relative to pulldown of BRD4-ET by the GST-only control (Figure 1C). A reciprocal experiment with immobilised GST-BRD4-ET domain and ASXL1^537-587^ fused to maltose binding protein also showed clear binding, indicating that ASXL1^537-587^ and BRD4 readily interact irrespective of direction of pulldown (Figure 1D).

Interactions between epigenetic regulators and ET domains have been characterised as ranging from relatively weak (7–950 *μ*M K_D_; (Wai *et al*, 2018)) to nanomolar (160 nM MLV; (Crowe *et al*, 2016)), albeit using various biophysical techniques. To compare ASXL1 binding-affinity in the context of previous interactions, we performed isothermal titration calorimetry with purified protein products. Titrating BRD4-ET into MBP-ASXL1^537-587^ we observed a dissociation constant of 535 nM, based on three replicate titrations (Figure 2A). Sub-*μ*M binding-affinity is also consistent with co-elution experiments, which showed that the same ASXL1 peptide epitope fused to lysozyme readily co-eluted with the BRD4-ET domain on size-exclusion chromatography (Supplementary Figure S1). Together, these analyses show that ASXL1 contains a functional motif for BRD4-ET-domain binding, which binds with relatively high affinity compared to other characterised ET-domain epitopes.

**Figure 2.**
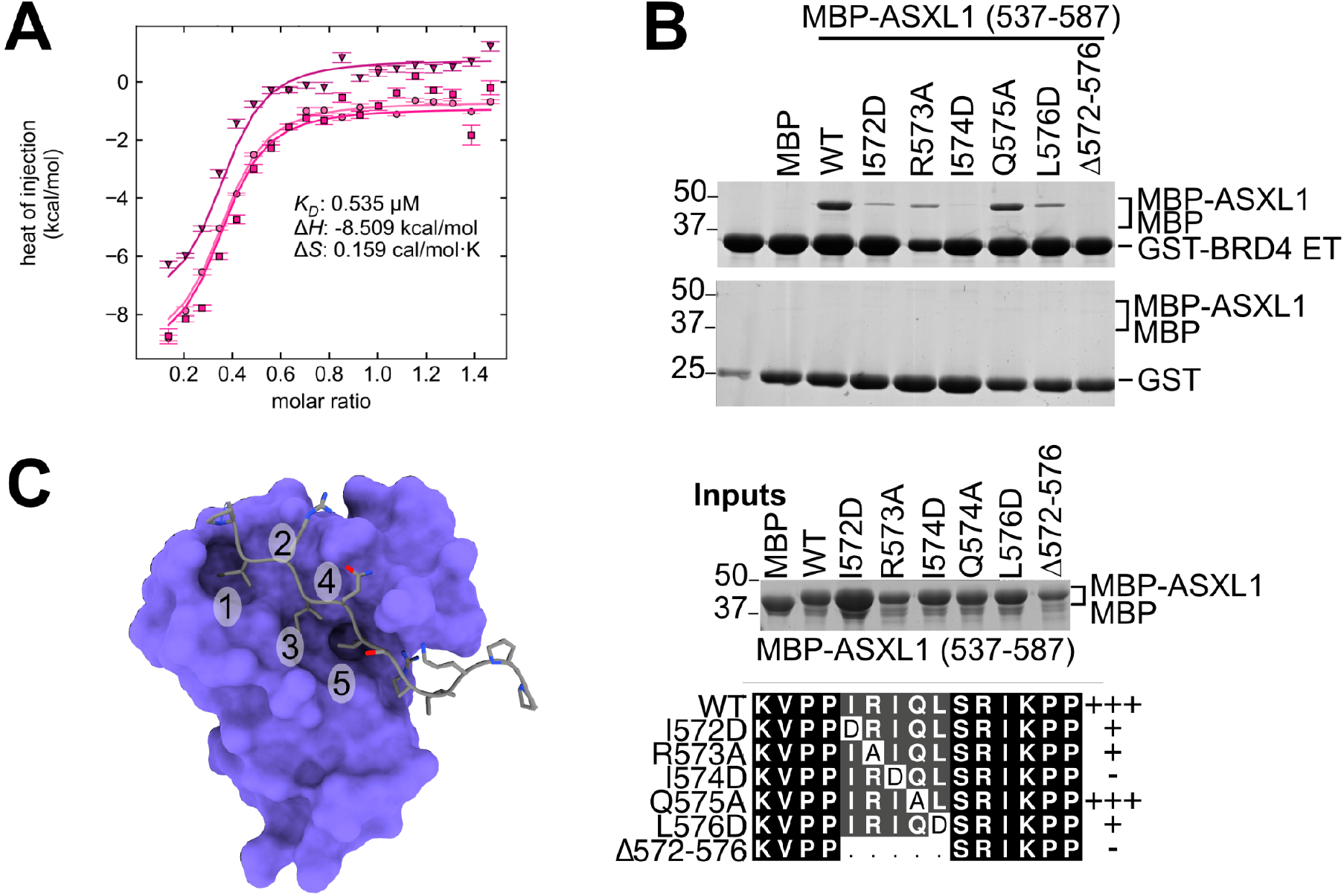
ASXL1 binds BRD4-ET through a core motif. (A) Isothermal titration calorimetry of ASXL1 and BRD4-ET. Isothermal titration calorimetry (ITC) of His_6_MBP-fused ASXL1^537-587^ injected into BRD4-ET domain reveals a high affinity interaction, with K_D_ 0.535 μM. Data points of triplicate titrations, following His_6_MBP titration subtraction. (B) Introduction of point mutations within the binding sequence of ASXL1 disrupts binding of His_6_MBP-ASXL1^537-587^ to GST-BRD4^600-678^. Coomassie-stained gels showing pulldown of His_6_MBP-ASXL1^537-587^ by GST-BRD4^600-678^. Multiple sequence alignment of the ASXL1 mutations below; the residues colored according to ALSCRIPT Calcons. (C) Model of ASXL1 binding to BRD4-ET (purple) based on PDB: 2NCZ; indicating the five residues of the core motif.

### Validation of the core BRD4-binding epitope in ASXL1

Multiple structures have been resolved using NMR spectroscopy of the ET domains of BRD4, and BRD3, in complex with binding partners (Zhang *et al*, 2016; Crowe *et al*, 2016; Wai *et al*, 2018). The ET-domain consists of three α-helices, a single β-strand and substrate proteins bind within a hydrophobic groove between two of the helices and the β strand. Substrates bind in such a manner that an antiparallel β-sheet is formed between the substrate and the ET domain. Interactions involving hydrophobic and basic residues within the ET-binding motif extend the interaction. The putative ET-binding motif in ASXL1 contains differences at several conserved residues within the canonical ET-binding motif—most notably arginine-573 where all characterised partners have a lysine, and glutamine-575 where all other interactors have an arginine or lysine (Figure 1B). Considering these differences, we validated that the putative motif was responsible for BRD4-binding and ascertain which residues might effect ASXL1-binding.

We created a series of mutations ranging from complete deletion of residues 572–576 (the entire core motif), to point mutants designed to disrupt the interaction if ASXL1 binds similarly to known binding partners of BRD4 (Figure 2B). Deletion of the core-motif completely abrogated pulldown of MBP-ASXL1^537-587^ by the BRD4-ET domain, validating that these five core residues of the motif are responsible for mediating interaction with the ET-domain. The individual point mutants disrupted the interaction to varying extents. Mutation of isoleucine to aspartate (I574D) was sufficient to disrupt binding alone, consistent with disruption of a hydrophobic residue at the centre of the motif. The mutations flanking this central position also reduced binding, with the greatest reduction in binding seen upon disruption of further hydrophobic residues (isoleucine-572 and leucine-576). Arginine-572, one of the ASXL1 residues that diverges from the generally conserved lysine, is also clearly important for the interaction based on reduced binding when mutated to alanine. The second divergent residue (glutamine-575) is apparently less important for high-affinity binding, given the Q575A mutant retains relatively avid binding.

The tightest measured ET-domain interactions are between BRD4 and MLV (160 nM; (Crowe *et al*, 2016)), and one of the two BRD4-binding sequences from NSD3 (2 μM; (Rahman *et al*, 2011)). To achieve tight binding MLV and NSD use the same core binding motif, but with either second β-strand (MLV) or with an extended interaction at the C-terminal end of the motif (NSD3; Supplementary Figure S2). To investigate whether high affinity ET-domain binding by ASXL1 requires residues beyond the core motif we made an expanded series of ASXL1 constructs to determine both the N- and C-terminal boundaries of the binding site. GST-pulldown assays showed that N-terminal truncation from residues 537– 567 were not critical for binding (Supplementary Figure S2A). Conversely, there was a decrease in co-precipitation of ASXL1 when residues between 577 and 587 to the C-terminus of the core motif was removed (Supplementary Figure S2B). Comparing the binding affinity of ASXL1^567-587^ with that of our initial construct ASXL1^537-587^ showed that these twenty residues are largely sufficient for binding, with only a two-fold decrease in the K_D_ for the longer construct (1.36 *μ*M; Supplementary Figure S2C). Together, these experiments strongly suggest that ASXL1 binds with relatively high-affinity by forming an extended single beta-strand in the manner of NSD3^152-163^, rather than the beta-hairpin formation seen for MLV (Supplementary Figure S2D; (Shen *et al*, 2015; Zhang *et al*, 2016; Crowe *et al*, 2016). Based on these results we created a model to visualise potential ASXL1-BRD4 binding based on NSD3 (Figure 2C). In this model, Arg578 aligns well with the basic lysine residue seen in an equivalent position of NSD3, consistent with a shared binding mode. Variation of residues within the ET-binding motif of ASXL1 also illustrates that more divergent sequences can bind to the ET domain, suggesting potential for a broader pool of ET-domain epitopes.

### Pairwise interactions are conserved across ASXL1–3 and BRD2–4

Both ASXL1 and BRD4 are part of multi-protein families with a conserved domain structure. An interaction between BRD4 and fulllength ASXL3 via the ET domain was reported while this work was in progress, which is proposed to modulate enhancer activity in lung cancer (Szczepanski *et al*, 2020). The authors reported no detectable interaction between ASXL2 or ASXL1 and BRD4. Nor were ASXL1/2 interactions reported in any other mass-spectrometry studies that we surveyed (Rahman *et al*, 2011; Mann *et al*, 2021; Dawson *et al*, 2011; Lambert *et al*, 2019; Szczepanski *et al*, 2020).

To test whether ASX-like proteins are more broadly capable of binding the ET-domains of bromodomain proteins, we first tested binding by the ET-domains from BRD2, BRD3 and BRD4 to ASXL1. Consistent with high-levels of sequence identity within the ET-domain, particularly surrounding the binding-groove, all three ET domains readily coprecipitated with ASXL1 (Figure 3A). Based on the ET-binding-motif in ASXL1, we aligned highly similar sequences in all three ASX-like proteins (Figure 3B). The most drastic variation was a proline in ASXL2 in the position corresponding to glutamine-575 in ASXL1, which we had already shown was tolerant to mutation (Figure 2B). We created MBP-fusion proteins of ASXL2^582-634^ and ASXL3^977-1027^ and saw that these fragments retained the same ability as ASXL1 to bind the GST-ET domain of BRD4 (Figure 3C). To ascertain if reports of different ASX-like protein binding by different family members could be explained simply differences in binding affinity driven by sequence differences in ASXL1–3 we quantitatively compared the interactions using ITC, titrating MBP-ASXL proteins into purified ET domain (Figure 3D). Calculated dissociation constants for these interactions were remarkably similar for all family members, with estimated K_D_s of 1.3, 1.1 and 2.0 *μ*M for ASXL1–3 respectively. Note that this value is approximately two-fold higher than that displayed for ASXL1 in Figure 2A, presumably because the titration was carried out in the opposite direction.

**Figure 3.**
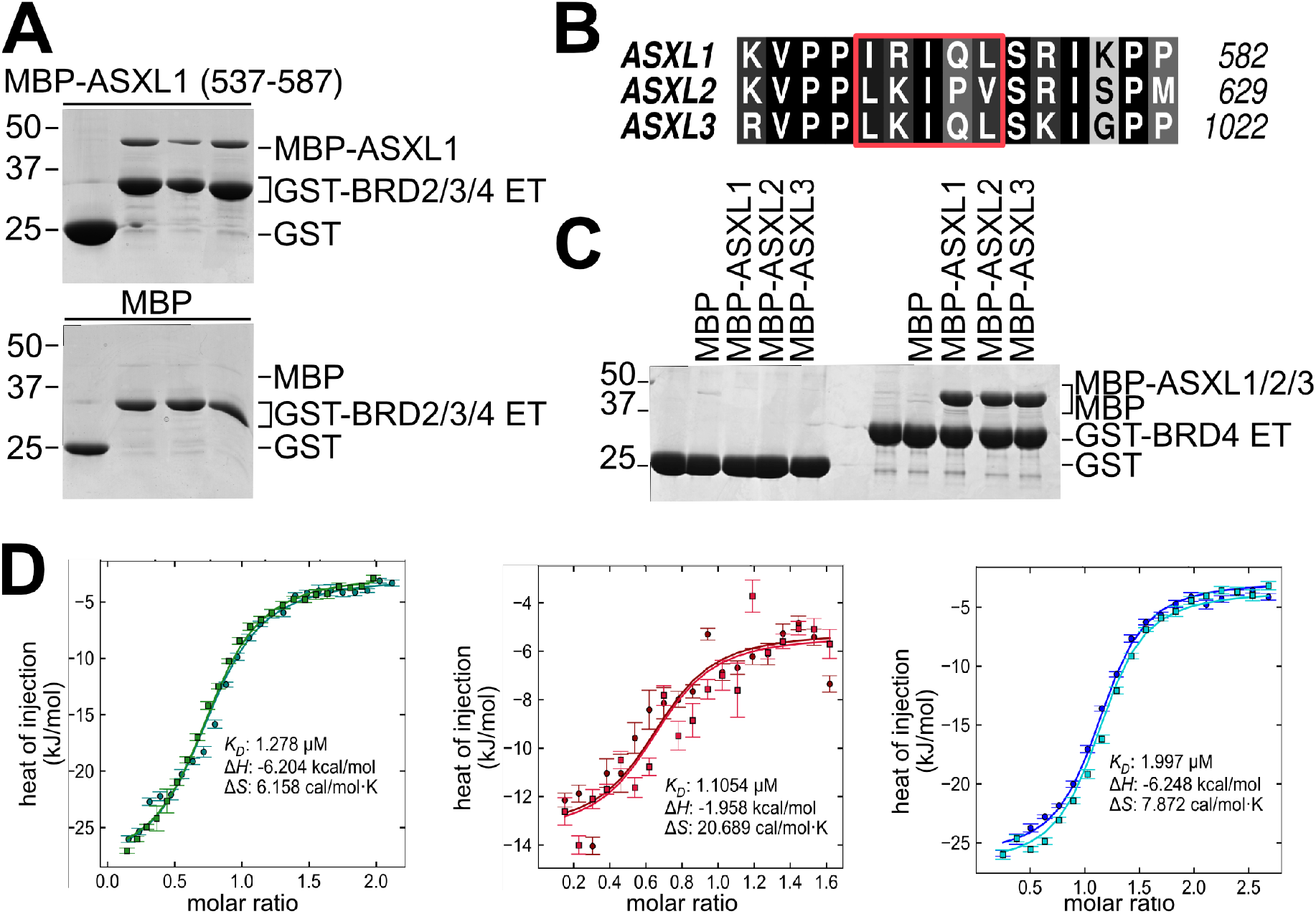
ASXL1-BRD4ET interaction is conserved. (A) ASXL1 binds to the highly conserved ET domain of BRD2, BRD3 and BRD4. Coomassie-stained gels showing pulldown of His_6_MBP-ASXL1^537-587^ by GST-BRD2^632-710^, GST-BRD3^562-640^ and GST-BRD4^600-678^. (B) Multiple sequence alignment of ASXL1, ASXL2 and ASXL3; the residues colored according to ALSCRIPT Calcons, demonstrates the conservation of the binding site; the core motif is highlighted (red box). (C) BRD4 binds ASXL1, ASXL2 and ASXL3. Coomassie-stained gels showing pulldown of His_6_MBP-ASXL1^537-587^, His_6_MBP-ASXL2^582-634^ and His_6_MBP-ASXL3^977-1027^ by GST-BRD4^600-678^. (D) Isothermal titration calorimetry of ASXL1 (green), ASXL2 (red) and ASXL3 (blue) injected into BRD4-ET domain reveals all ASXL homologues tightly bind the BRD4-ET domain. Data points of triplicate titrations, following His_6_MBP titration subtraction.

Together these results show an ET-binding motif is conserved amongst ASX-like proteins, and able to bind across BRD2–4. The similarity of binding-affinity for the ET-domain across different ASX-like proteins suggests that differential binding by full-length proteins is unlikely to be due to any inherent inability to bind. Rather the availability of the binding sites is likely to vary in different cell types or contexts.

### Truncation of ASXL1 enhances BRD4-ET binding

We next sought to discover whether truncated ASXL1 variants expose an ET-binding motif that is non-functional in full-length protein, or if truncation merely enhances a normally weak interaction that is difficult to detect by co-immunoprecipitation. Where various studies have elegantly analysed the broader impact of ASXL1 truncating variants on epigenetic histone marks, a more targeted system was required to measure the difference in BRD4-binding— and hence potential regulatory crosstalk—between different lengths of ASXL1 truncation. To achieve the sensitivity and quantitation required, we applied a variant of proximity ligation by TurboID (Branon *et al*, 2018), rationalising that its sensitivity would allow detection of weak or transient interactions and facilitate downstream quantitative mass spectrometry. HEK293T cells were transiently transfected with various constructs of ASXL1, and the BRD4-ET domain fused to the TurboID biotin ligase (Figure 4A). When treated with biotin, proteins which are associated with/or are in close-proximity of the BRD4ET domain are biotinylated, and as such can be detected and/or enriched by streptavidin affinity purification.

**Figure 4.**
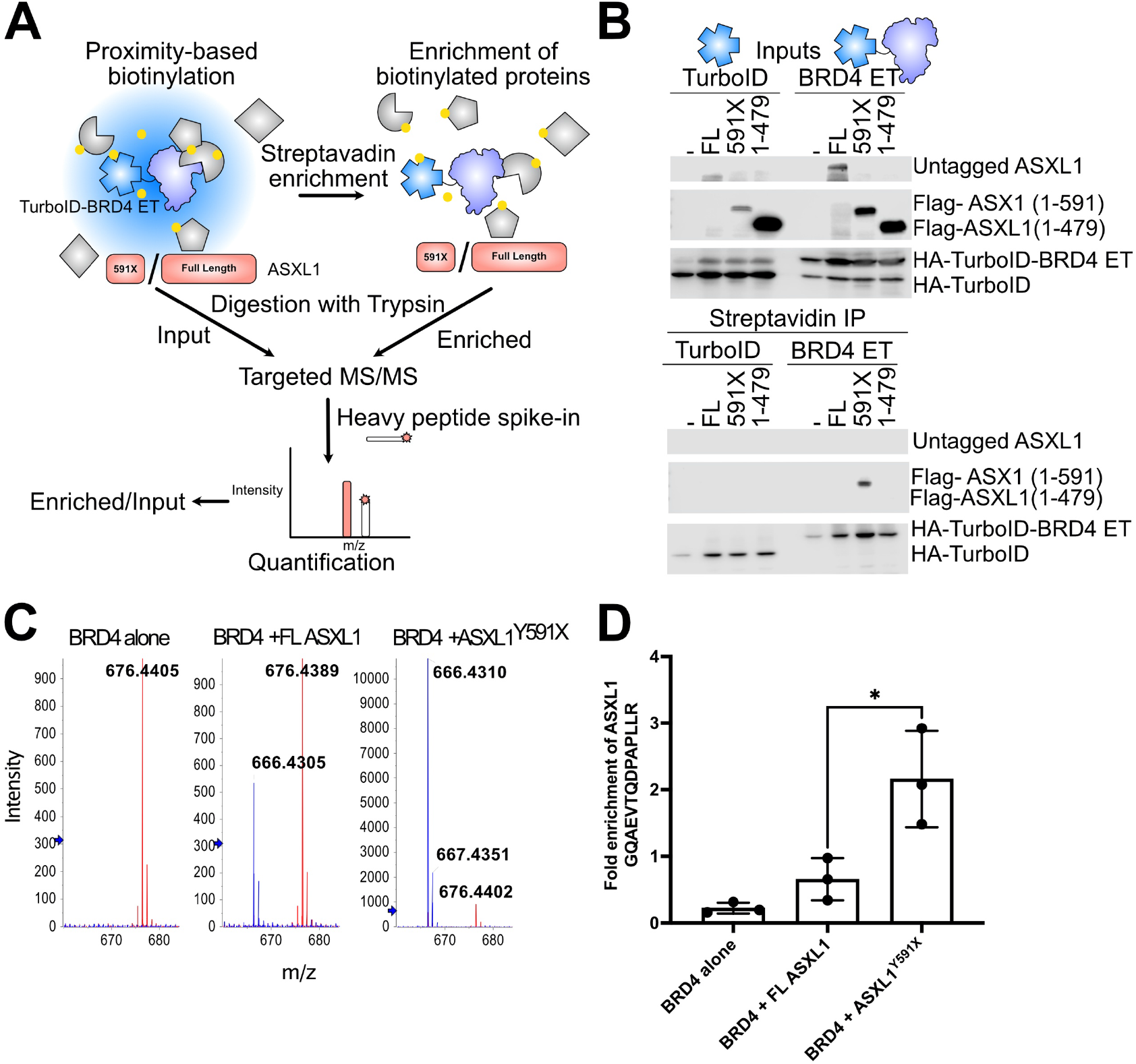
Enhanced BRD4-ET binding by ASXL1^Y591X^. (A) Schematic showing the approach used for targeted analysis. TurboID was fused to BRD4-ET and used to investigate the interaction with different lengths of ASXL1 (B) Proximity based labelling assays indicate that BRD4 interacts with ASXL1^Y591X^ and neither the full-length, nor ASXL1^1-472^ which lacks the ET binding site. HEK293T cells were transiently co-transfected with either TurboID-BRD4 or TurboID alone and varying lengths of ASXL1. Cells were treated with biotin for ten minutes prior to lysis. Immunoprecipitation and Western blotting were performed using to determine if ASXL1 was biotinylated. (C) Quantification of ASXL1 enrichment using parallel reaction-monitoring (PRM) mass spectrometry with isotopically-labelled peptide to determine fold enrichment of ASXL1 after one hour treatment with biotin and immunoprecipitation. (D) Representative ion intensities of the strongest y-ion (y6) of the GQAEVTQDPAPLLR peptide in overlaid spectra of the endogenous peptide (blue trace) and heavy stable isotope labelled internal standard peptide (red trace).

We first tested: full-length ASXL1; the commonly occurring ASXL1^Y591X^ truncation; and ASXL1^1-479^, which contains the HARE-HTH and Deubad domains of ASXL1 but not the ET-binding motif. Analysing this experiment by western blotting, all three of the ASXL proteins could be detected to varying levels in input samples, indicating that they were expressed. However, only ASXL1^Y591X^ could be detected following affinity purification with streptavidin, providing qualitative data that the BRD4-ET domain preferentially interacts with the Y591X truncated protein, but not the full-length protein, or ASXL1^1-479^ which lacks the ET-binding motif (Figure 4B). The difference between proteins ending at 591 and 479 validates that the core ET-binding motif is active in cells, but expression levels make it difficult to compare relative association between the truncated and full-length proteins.

To account for varying expression levels, we employed quantitative assessment of ASXL1 input and affinity-purified proteins using parallel reaction-monitoring (PRM) mass spectrometry and stable isotope-labelled internal standard (SIS) peptides for sequences from the N-terminal portion of ASXL1. Here we could directly compare the relative abundance of ASXL1 in input samples, with samples that have been biotinylated through association with BRD4-ET domain. To increase sensitivity and enable detection of very weak association with full-length ASXL1, we increased the biotinylation time of the experiment to 1 hour, whereas western-blotting was performed after only a 10 minute biotin treatment. Normalising the amounts of ASXL peptides between input and streptavidin-enriched samples, we observed approximately three-fold enhanced ET-domain binding by ASXL1^Y591X^, compared with fulllength ASXL1 (Figure 4C/D, Supplementary S3C). The same analysis using a second ASXL1 peptide showed approximately six-fold enhanced ET-binding but was more variable (Supplementary Figure S3). These data show that ET-domain binding can occur to a limited extent in the context of full-length ASXL1 but is several-fold enhanced in the context of truncated protein.

### ASXL1 truncation variants recruit BRD4 to relevant regulatory complexes

To assess if the differences in ASXL1-ET binding manifest as enhanced recruitment to the PR-DUB, or a change in the complexes where BRD4 is recruited, we analysed the global profile of BRD4-ET domain interactors following streptavidin enrichment. As expected based on the targeted analysis (Figure 4), comparing transfection of full-length ASXL1, or ASXL1^Y591X^, to cells transfected only with TurboID-BRD4-ET showed that ASXL1 was significantly enriched for either ASXL1 construct (Figure 5A/B). A number of other proteins that have previously been identified to interact with the BRD4-ET domain, CHD1, CHD4 and BRG1, were also detected following streptavidin enrichment (Supplementary Figures S4 and S5; (Rahman *et al*, 2011; Wu *et al*, 2018; Conrad *et al*, 2017)). While the enrichment of neither BAP1 nor HCFC1 was statistically significant with full-length ASXL1, BAP1 was significantly enriched with ASXL1^Y591X^ (Figure 5, Supplementary Figure S5). This indicates that the BRD4-ET domain is specifically recruited to biologically-relevant ASXL1-containing complexes, but to different extents. To finally assess if there were differences in the specific types of PR-DUB complexes that BRD4-ET is recruited to, we compared the BRD4-ET interactomes when it was transfected with different lengths of ASXL1 (Figure 5C). Notable from this analysis was: the PHD finger protein PHF10, which is enriched in the presence of ASXL1^Y591X^ but not the full-length protein; and the MAZ transcription factor, that was strongly enriched as a BRD4-ET interactor with either length of ASXL1. To iterate that these are likely to be specific interactions, we also analysed the interactome profile against commonly co-purified contaminating proteins (Mellacheruvu *et al*, 2013), which showed that ASXL1, MAZ and PHF10 are infrequently co-purified in unrelated co-IP experiments (Supplementary Figure S5). Together, these results illustrate that enhanced interaction between ASXL1 and BRD4 has the potential to mis-direct epigenetic crosstalk of the two proteins in a reciprocal manner. First, that co-localisation of BRD4 with different epigenetic regulators such as ASXL1 and PHF10 may be different depending on the length of ASXL1 protein present in the cell. And secondly, that interaction of the BRD4-ET domain with ASXL1 appears to strongly upregulate its co-localisation with the MAZ transcription factor, relative to the BRD4-ET domain alone.

**Figure 5.**
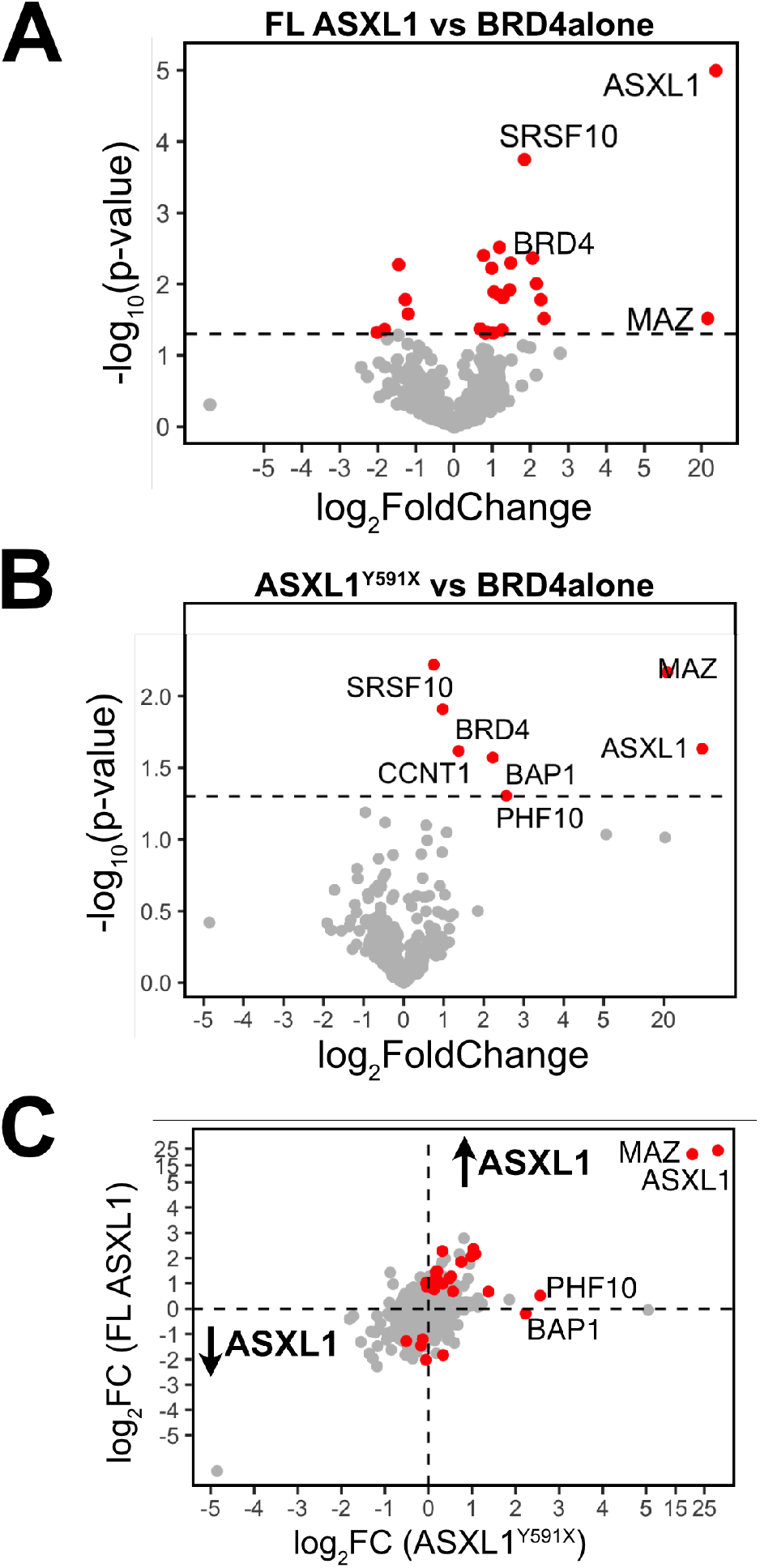
Global Protein Profile Analysis of ASXL1 lengths. Protein identification following streptavidin enrichment of cells treated with TurboID-BRD4 alone or TurboID-BRD4 with either Full-length ASXL1 or ASXL1^Y591X^. A volcano plot was generated to demonstrate changes between proteins detected with FL ASXL1 (A) or ASXL1^Y591X^ (B) compared with the control (BRD4 alone). The fold-change (ASXL1/BRD4 alone) is plotted against the statistical significance (-log10p-value). Horizontal dashed line indicates where p = 0.05/ Proteins with significant p-values are highlighted (red) and labelled. **(C)** Comparative plot of FL ASXL1 and ASXL1^Y591X^. The fold-change (ASXL1 FL/BRD4 alone) is plotted against the fold-change (ASXL1^Y591X^ /BRD4 alone). Proteins which are significantly changed in either data set are highlighted (red).

## DISCUSSION

Here we delineate the basis for BRD4 recruitment by ASXL proteins, and show that this interaction is significantly enhanced in the context of a common truncation variant of ASXL1 that occurs in acute myeloid leukaemia patients. This work provides a detailed molecular basis for the finding of Yang et al, who first reported that truncation of ASXL1 generates a new interaction with BRD4 that does not occur in the full-length protein (Yang *et al*, 2018). While previous co-immunoprecipitation studies suggested multiple possible interaction sites, our work shows that a peptidic motif comprised within ASXL1 is necessary and sufficient to bind to the BRD ET domain. This interaction is consistent with ASXL1 binding as an extended β-strand in the manner of other ET-interactors, and is relatively high-affinity compared to other ET-interactors. By using quantitative targeted mass-spectrometry we could control for different protein stabilities of full-length and truncated ASXL1, show that BRD4-ET binding occurs to some extent in the context of full-length ASXL1 but is 3.5-fold enhanced in the context of ASXL1^Y591X^ truncation. These results support a model where a ASXL1 has a limited ability to bind the ET domains of BRD proteins in a native context, but truncating mutations release a constraint on binding and enhance BRD4 recruitment. Thus, increased binding propensity between truncated and full-length ASXL1, could combine with potential protein stability effects, to enhance co-localisation and epigenetic co-regulation between BRD4 and ASXL1.

The clear next question arising from this model is how truncating mutations precisely enhance recruitment of ET domains. We can envisage two possible mechanisms for how binding-enhancement may occur. Either truncation could alter local secondary structure around the ET-binding motif, and so create a protein with a more optimal binding structure for ET-domains; or the ET-binding motifs are normally allosterically blocked by other binding partners in the context of full-length protein, which are lost in the context of truncated proteins. Since the initial report of BRD4 binding by truncated ASXL1, Szczepanski *et al* (2020) have reported that full-length ASXL3 can bind to BRD4 whereas full-length ASXL1 or ASXL2 cannot. Our survey of previous ASXL1, and BRD protein proteomic studies also suggest that BRD proteins interacting with ASXL1 and ASXL2 is infrequent or undetectable in most circumstances. However, targeted mass spectrometrybased assay suggests that binding in the context of full-length ASXL1 occurs to a limited extent but is below the limit of detection of western blotting. Our in vitro data suggests that the minimal ET-binding motif of ASXL3 binds with to the ET domain with a similar affinity as the motifs from ASXL1 and ASXL2, reiterating that other factors likely regulate association in the cellular context. Overall, this supports a model of steric restriction of BRD4 binding the full-length proteins under most circumstances, which is relieved after truncation. Either incorporation into specific protein complexes, or post-translational modifications, could impede ET-binding motif of full-length ASXL1 and ASXL2 under normal circumstances, but these factors are released when ASXL1 is truncated.

Further clues as to the mechanisms governing ASXL–BRD protein binding under normal circumstances may come from the finding that full-length ASXL3 can detectably bind to BRD4. Where ASXL1/2 are more globally expressed, ASXL3 is predominantly expressed in the brain and pluripotent respiratory epithelial cells (Shukla *et al*, 2017). It is possible that the factors that sterically hinder ET-recruitment may be less of a factor in these tissues, but further analysis of ASXL interactors will be required to understand if ET-recruitment is a key functional difference between full-length ASXL proteins. It is relevant to note that ASXL3 truncating mutants also occur in Bainbridge–Ropers syndrome (Srivastava *et al*, 2016). It may be relevant to understand whether the presence or absence of the ET-binding motif could affect presentation or penetrance of the condition, or potentially inform therapy in the future.

Perhaps the best know role of ASXL1-3 is in binding and activating the polycomb-repressive deubiquitinase, BAP1. ASXL1–3 have been reported to form mutually exclusive complexes with BAP1, and of particular functional relevance truncated ASXL1 is stabilised in a mono-ubiquinated form, whereas the full-length protein is subject to polyubiquitination proteasomal degradation (Daou *et al*, 2018; Asada *et al*, 2018). The BRD4 recruitment we characterise here may either be an active player in the regulated turnover of ASXL1 full-length protein; or be an additive factor. For instance, avid recruitment of BRD4 may scaffold complexes that contain deubiquitinases that act on truncated ASXL1, although we did not identify particular deubiquitinase candidates apart from BAP1 itself. In addition, truncated ASXL1 could redirect BRD4-containing complexes to drive aberrant gene expression. Our finding that ASXL1 drives BRD4 co-localization with the MAZ transcription could be particularly relevant in this regard. For instance, MAZ has: a known role co-operating with MYC to regulate transcription (Bossone *et al*, 1992), and disrupted MYC-regulated gene expression was a signature of ASXL1-truncated mice (Yang *et al*, 2018; Zhang *et al*, 2018); an SP1-like DNA-binding specificity that is similar to that previously observed for the PR-DUB (Dey *et al*, 2012); and an ability to interact with the nucleosome-remodelling factor subunit BPTF (Jordan-Sciutto *et al*, 2000).

Whatever the precise mechanisms at play, ASXL1-BRD4 binding could be relevant to two different small-molecule based strategies in ASXL1 truncated cancers. Firstly, the BET-domain antagonist JQ1 can modulate PR-DUB regulated gene expression in ASXL1^Y588X^ mice (Yang *et al*, 2018). Secondly, it has recently been shown that BAP1 inhibitors can also impair leukaemia progression in the context of ASXL1 gain-of-function truncations (Wang *et al*, 2021). The ASXL1–3-BRD4 interaction characterised here provides a mechanistic basis for crosstalk between these two proteins, and suggest there may be benefit in testing the combined effects of the BET domain inhibition and BAP1 inhibition in the future.

## METHODS

### Plasmids and cloning

DNA constructs for recombinant protein expression in *Escherichia coli* were synthesised by Integrated DNA Technology (IDT), with the 5’-(CAGGGACCCGGT) and 3’-(TAACCGGGCTTCTCCTCG) overhangs, required for Ligation-Independent Cloning (LIC). Constructs were cloned into a pET-LIC vector either containing an N-terminal His_6_ and His_6_MBP or GST tag and a 3C protease cleavage site.

DNA constructs for protein expression in HEK293T cells were produced as follows. ASXL1 truncations were purchased from Addgene: pCMV6-XL4 ASXL1 (1-1304) (Addgene # 74244); pCMV6-XL4 ASXL1 (p.Y591X) 3x FLAG (Addgene # 74261); pCMV6-XL4 ASXL1 (1-479) 3x FLAG (Addgene # 74262) were a gift from Anjana Rao. Full-length ASXL1 in pcDNA3.1 was synthesised by Genscript. The BRD4-ET domain (600-678) was cloned into pcDNA3 containing TurboID modified for LIC cloning. This vector was modified from 3xHA-TurboID-NLS_pCDNA3 which was a gift from Alice Ting (Addgene # 107171).

### Expression and Purification of Recombinant Protein

The BRDET domains and ASXL Peptides were expressed in *Escherichia coli* BL21 (DE3) cells. Cells were grown at 37 °C in either Terrific broth (TB) or Luria-Bertani (LB) medium to an OD_600_ of approximately 0.6, then induced with 0.2 mM isopropyl-β-D-1thiogalactopyranoside (IPTG) and grown shaking at 18 °C overnight (O/N).

Cells were lysed by sonication in a buffer containing 50 mM Tris-HCl (pH 8.0), 300 mM NaCl, 10 % (v/v) glycerol, 10 % (w/v) sucrose, 10 mM imidazole, and supplemented with DNase I (AppliChem) and Lysozyme (0.2 mg/mL). The soluble fraction was bound to the appropriate resin, either Ni^2+^-nitrilotriacetic acid (NTA) resin or GSH resin and washed three times in the same buffer. Protein was eluted from the Ni resin by increasing the concentration to 300 mM imidazole. Protein immobilized of GSH resin was retained on the resin for pull down experiments.

### Isothermal Titration Calorimetry (ITC)

BRD4-ET and MBP fusion proteins of ASXL1 (537-587), (567-587), ASXL2 (582-634) and ASXL3 (977-1027) were initially purified by Ni^2+^ affinity chromatography, using a 5 mL HisTrap FF column (GE Healthcare). Bacterial pellets were lysed in a buffer containing 50 mM Tris pH 8.0, 300mM NaCl, 10 mM Imidazole, 10% Sucrose and 10% Glycerol, and the supernatant was loaded onto a 5 mL HisTrap column (GE Healthcare); which was then eluted using a step to increase the concertation of imidazole to 300 mM. Fractions containing the purified protein were recovered. The His_6_ tag was removed by 3C protease digestion at 4 °C overnight, with 2 mM dithiothreitol (DTT). Protein was further purified on a Superdex 75 *Increase 10/300* GL or Superdex 200 Increase 10/300 GL, as appropriate (GE Healthcare) in a buffer containing 20 mM HEPES pH 7.6 and 300 mM NaCl. Proteins were then dialysed against this same buffer overnight at 4 °C to ensure buffer matching. ITC was performed by titrating BRD4-ET into MBP-ASXL1 and MBP or the reciprocal. ITC with ASXL2 and ASXL3 were performed by titrating MBP-ASXL constructs into BRD4-ET. The titration was performed using a MicroCal VP-ITC at 25 °C with 30 injections (2.5 *μ*L injections) with 200s between injections. Baseline corrections and integration were performed using NITPIC, isotherms were fit to a single site-binding model using SEDPHAT, and figures were generated using GUSSI (http://biophysics.swmed.edu/MBR/software.html).

### Immunoprecipitation

Proteins immobilised on GST Sepharose resin (Amersham Biosciences) were analysed for purity by SDS-PAGE. GST-ASXL1 (537-587) was incubated with His_6_BRD4 (600-678); GST-BRD2 (632-710), GST-BRD3 (562-640) and GST-BRD4 (600-678) were incubated with varying lengths of His_6_MBP-ASXL1, His_6_MBP-ASXL2 (582-634) and His_6_MBP-ASXL3 (977-1027) at 4 °C for 20 min using phosphate-buffered saline (PBS; pH 7.4) with 0.02 % (v/v) Tween 20 and 2 mM DTT. After incubation, the resins were washed three times in this buffer. Samples were then resuspended in SDS sample buffer and visualized on 12-18 % gradient SDS-PAGE stained with Coomassie R250.

### Analytical Size Exclusion

The elution profile of ASXL1(537-587) fused to lysozyme and BRD4-ET either alone or together was evaluated by injecting each purified protein onto a Superdex 75 *Increase 10/300* GL Increase column (GE Healthcare) equilibrated with 20 mM HEPES 7.5, 300 mM NaCl, 2 mM DTT and the absorbance monitored at 215 nm. The lysozyme fusion created a larger size discrepancy between the two separate proteins to ensure complex formation could be determined. The BRD4-ET used in these experiments was purified as a GST fusion and then the tag removed prior to analytical size exclusion.

### Cell Lines and Cell Culture

HEK293T cells were grown in Dulbecco’s modified Eagle’s medium (DMEM; Life Technologies, 10566) supplemented with 10 % fetal bovine serum (Sigma-Aldrich, F8067), 2 mM L-glutamine (Life Technologies, 25030081), Penicillin-Streptomycin (100 U/ml; Life Technologies, 15140122), Non-Essential Amino Acids (HyClone, SH30238), and 1 mM Sodium Pyruvate (HyClone, SH3023901). Cells were grown at 37 °C in a humidified atmosphere with 5 % CO_2_.

#### Biotin treatment and streptavidin affinity purification

HEK293T cells were transiently co-transfected with Lipofectamine 3000 (Life Technologies, L3000015) to express either TurboID or TurboID fused to BRD4-ET and/or varying lengths of ASXL1. Cells were grown for forty-eight hours before biotin treatment. Cells were treated with either DMSO (vehicle control) or 50 *μ*M biotin for 10 mins in the initial assay or 1 h for targeted analysis. Cells were harvested into the treatment medium and centrifuged at 1000 x g for 5 min at 4 °C. The cell pellets were washed four times with ice-cold phosphate-buffered saline (PBS; pH 7.4). Cell pellets were frozen in liquid nitrogen and stored at −80 °C until use. Before use cell pellets were thawed and resuspended in radioimmunoprecipitation assay buffer (RIPA buffer).

Lysates from treated HEK293T, were incubated with streptavidin beads for 1 hour at room temperature and then incubated overnight at 4 °C in rotation. The beads were washed four times with PBS with 0.02 % (v/v) Tween 20 to remove non-biotinylated proteins. Samples were then either resuspended in 4× SDS–polyacrylamide gel electrophoresis (SDS-PAGE) sample buffer and analysed by western blotting or used for mass spectrometry analysis.

Full-length ASXL1 has been reported to have limited activity and expression when tagged; to avoid any compounding difficulties in binding that may result from this, we used an untagged protein. Truncations of ASXL1 were N-term FLAG tagged.

### Western Blotting

For analysis by western blot, samples were separated by SDS/PAGE and transferred to 0.2 *μ*m nitrocellulose (Life Technologies, IB23002). Membranes were blocked in 5 % BSA (w/v) in TBS-T. Membranes were incubated with primary antibodies overnight at 4 °C in 1 % BSA (w/v) in TBS-T. Antibodies used in this study were: mouse monoclonal α-HA (F-7) (1:1000, sc-7392), mouse monoclonal α-ASXL1 (1:1000, ab50817, C-terminal epitope) and/or rabbit monoclonal DYKDDDDK tag (Flag tag, 1:1000, CST, #14793). Following three washes with TBS-T, membranes were incubated with secondary antibodies diluted in TBS-T with 1 % (w/v) BSA for 1 hour at room temperature. Secondary antibodies used were goat anti-rabbit IRdye 680LT (LI-COR), goat anti-mouse IRdye 800LT (LI-COR) or goat anti-rabbit HRP-conjugated (1:10,000, Abcam, ab6721). Membranes were washed a further three times with TBS-T. HRP-conjugated secondary antibodies were detected using Clarity Western ECL Substrate (Bio-Rad). Membranes were then developed with the Odyssey Fc imaging system.

### TurboID and Mass Spectrometry Analysis

To determine relative intensities of full-length and truncated ASXL1 that interact with BRD4 ASXL1 was co-transfection with TurboID-BRD4-ET as described above. Following one hour of treatment with biotin cells were harvested and lysed in RIPA buffer. Twenty percent of each lysate was removed and kept as an input sample to measure the endogenous amount of ASXL1. The remaining 80 % was bound to streptavidin beads. Beads were washed three times with RIPA buffer and then twice with PBS. The comparison of BRD4 alone, BRD4+ FL ASXL1 and BRD4+ ASXL1^Y591X^ was performed in three biological replicates, i.e. three independent transfections of each sample and independent processing as well as measurements of each set of samples.

Pull-down samples were on-bead digested with trypsin. Therefore, proteins bound to the beads were washed twice with 200 *μ*l of 100 mM aqueous triethylammonium bicarbonate (TEAB), reduced and alkylated with 5 mM tris(2-carboxyethyl)phosphine (TCEP) and 20 mM iodoacetamide respectively and washed again twice with 200 *μ*l of 100 mM aqueous TEAB before incubating with 2 *μ*g of trypsin (proteomics grade, Promega) at 37°C for 4h. Samples were then boosted with 1 *μ*g of trypsin and digested at 37 °C for an additional 14 h. The supernatants were recovered by centrifugation at 20,000 x g for 30 min and dried using a centrifugal vacuum concentrator.

Input samples were processed and digested using the S-Trap method following the manufacturer’s protocol (ProtiFi, Farmingdale, NY) and subsequently dried using a centrifugal vacuum concentrator.

#### Parallel reaction monitoring mass spectrometry

To determine the molar ratio of ASXL1 before and after immunoprecipitation, we used two stable isotope labelled internal standard (SIS) peptides from the N-terminal region of ASXL1 (GQAEVTQDPAPLLR and IQAEPDNLAR) for absolute quantification of both the truncated and full-length form by parallel reaction monitoring (PRM) mass spectrometry. Protein digest of the pull-down and input samples were reconstituted in 20 *μ*l of 5 % acetonitrile in 0.1 % aqueous formic acid and split up in two 10 *μ*l aliquots, one for targeted PRM analysis and the other for shotgun proteomics. The 10 *μ*l aliquots for PRM analysis were mixed with 30 *μ*l of 5 % acetonitrile in 0.1 % aqueous formic acid containing 200 *ƒ*mol /*μ*l of each of the SIS peptides. 5 *μ*l (containing 25 *ƒ*mol of each SIS peptide) of each sample were loaded onto an in-house packed emitter tip column packed with Synergi^™^ 4μm Hydro C18 beads (Phenomenex) on a length of 20 cm. Peptides were separated through a 15 min linear gradient between 5 and 40 % (v/v) acetonitrile in 0.1 % (v/v) aqueous formic acid at a flow rate of 1000 nL/min using and Ekspert 415 nano LC-system (Eksigent; AB Sciex) inline coupled to the nano-spray source of a 5600+ TripleTOF mass spectrometer (AB Sciex). The mass spectrometer was monitoring fragment ion spectra of the calculated m/z values of 563.799, 568.802, 747.902 and 752.902 for the doubly charged precursor ions of light (endogenous) and heavy (SIS) IQAEPDNLAR and GQAEVTQDPAPLLR peptides respectively over the full-length of the gradient. A quadrupole precursor ion isolation window of 0.7 m/z and an ion accumulation time of 150 ms for each of the four fragment ion spectra was selected, resulting in a total cycle time of 1000 ms including a 350 ms survey scan. The collision energy was set to rolling. Each sample was measured in three technical replicates to estimate the intra-assay variation using the coefficient of variation in percentage (CV %).

#### Quantification of ASXL1

The skyline software (https://skyline.ms/project/home/begin.view) was used for quantification of endogenous and SIS peptides using the peak intensities of the 4 strongest y-fragment ions (containing the light- or heavy-isotope containing C-terminal arginine) above m/z 400. The absolute amount of endogenous full-length and truncated ASXL1 in each sample was calculated based on the y-fragment ion intensities of endogenous versus SIS peptide considering the total sample volume and dilution factors. The calculated amounts were averaged over the three technical replicates of each sample and the CV % was calculated. The amount of ASXL1 detected in each output sample was then directly compared to the corresponding input sample to determine the enrichment of ASXL1 following streptavidin affinity purification. The overall CV % was less variable with GQAEVTQDPAPLLR, and so this was primarily used for quantification (Supplementary S3). The samples which did not contain ASXL1 had a much higher CV %, the peaks themselves are broad-nonspecific- and do not align with the ASXL1 standard.

Analyses to determine significance between the biological replicates using t-tests and descriptive statistics were performed using Prism 9 (GraphPad Software). Due to the nature of samples provided, the input and output required different processing prior to quantification. While input samples were in-solution, the output samples were bound to beads. This difference in processing means that each sample treatment was normalised to the control sample treated in an equivalent manner, and fold-enrichment calculated relative to normalised control.

#### Global Protein Profiling

The remaining 10 *μ*l aliquots of each output sample from all three biological replicates were used for unbiased shotgun proteomics to identify all proteins that were detectable after streptavidin-biotin affinity enrichment. Therefore, samples were subjected to liquid chromatography coupled tandem mass spectrometry (LC-MS/MS) using an Ultimate 3000 RS LC-system inline coupled to the nano-spray source of an LTQ-Orbitrap XL mass spectrometer (Thermo Scientific). The total ion counts of the MS1 signal intensities from the PRM analysis were used to calculate normalization factors for equal sample loading. After adjustment of sample volumes 5 *μ*l of each sample were loaded onto an in-house packed emitter tip column filled with Aeris^™^ 2.6 *μ*m Peptide XB-C18 beads (Phenomenex) on a length of 20 cm. Peptides were separated through a 70 min linear gradient between 5 and 45 % (v/v) acetonitrile in 0.1 % (v/v) aqueous formic acid at a flow rate of 300 nL/min. The orbitrap analyser was operated at a resolution of 100,000 for precursor ion scans in a mass range of m/z 400 – 2000. The ten strongest precursor ion signals were selected for collision induced dissociation (CID) MS in the ion trap analyser allowing for 2 repeat acquisitions of CID spectra per precursor during a time window of 90 s. Raw data were processed through the Proteome Discoverer software (version 2.4; Thermo Scientific) and searched against the human reference sequence database (https://www.ncbi.nlm.nih.gov/refseq/, 87,570 sequence entries) using the Sequest HT search engine. A significant protein/protein group identification required at least two peptide hits with a significant score at a false discovery rate (FDR) of 1 % calculated by the Percolator algorithm (Käll *et al*, 2007). Precursor ion intensities were used for protein quantification using the top3 approach (Silva *et al*, 2006). The top three intensities were extracted and further analysed in RStudio to calculate fold changes and statistical significance (p ≤ 0.05). Any protein which was not detected in all three of the biological replicates of any single condition; BRD4 alone, BRD4+ FL ASXL1 or BRD4+ ASXL1^Y591X^ was given an arbitrary intensity of 1. Anything which was only detected in one of three biological replicates was not considered for comparison.

To visualise changes in protein enrichment each ASXL1 containing sample was compared to BRD4 alone. The mean peptide intensity of BRD4 + ASXL1 was divided by that of BRD4 alone to produce a fold-change (FC). The log_2_FC was then plotted against the −log_10_p-value and anything with a p-value ≤ 0.05 was highlighted in red on the volcano plots.

The total list of proteins detected in this analysis was submitted to the crapome (Mellacheruvu *et al*, 2013). This produced a list of how many times each protein occurs in unrelated studies and so each protein was assigned a score based on this = occurrence/ total number of studies. This was defined as the crap_score. The log_2_FC of each ASXL1^Y591X^ and FL ASXL1 (against BRD4 alone) was then plotted against the crap_score. Proteins of interest are those which have a low crap_score but are significantly changed (highlighted in red).

Figures were generated using RStudio (*http://www.rstudio.com/*.)

PDB files used:2NCZ, 2N3K, 6BGG

---

**Supplementary Figure S1.**
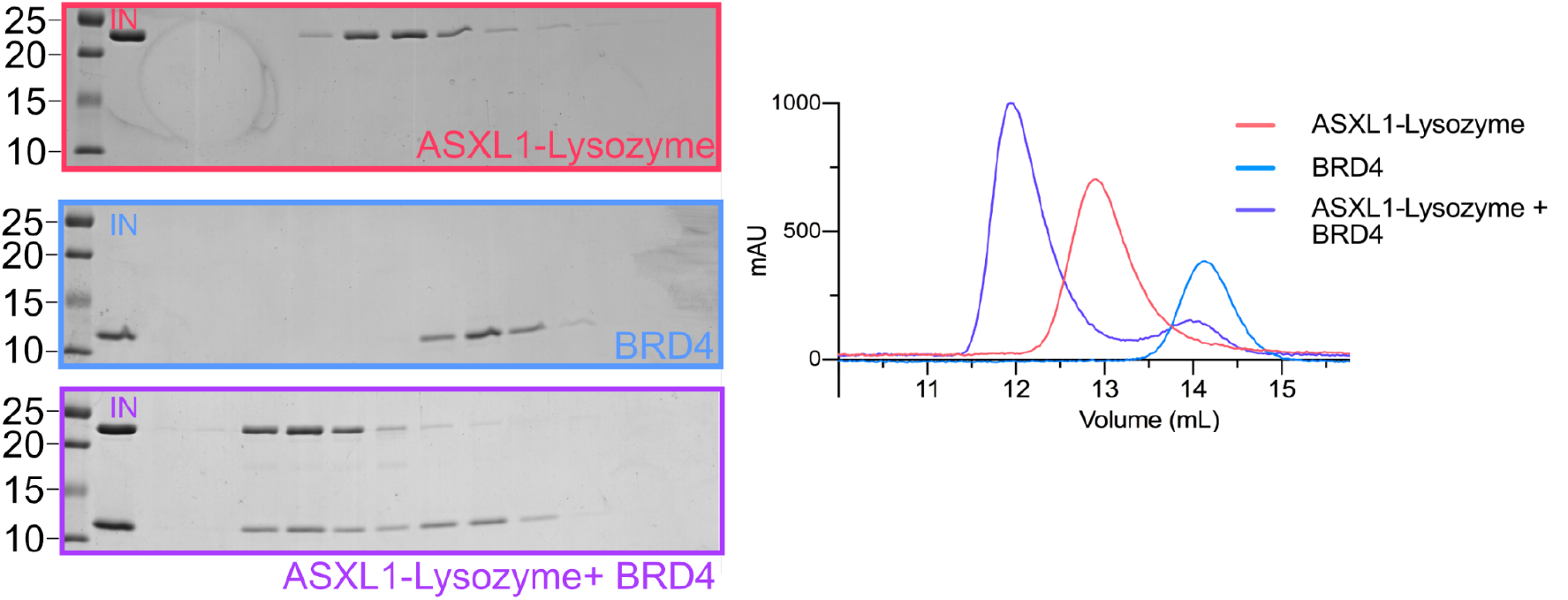
ASXL1-BRD4 Complex formation. Analytical size exclusion chromatography (SEC) of His_6_-ASXL1^567-587^-Lysozyme (Red) and His_6_BRD4^600-678^ (Pink), showing complex formation (Purple).

**Supplementary Figure S2.**
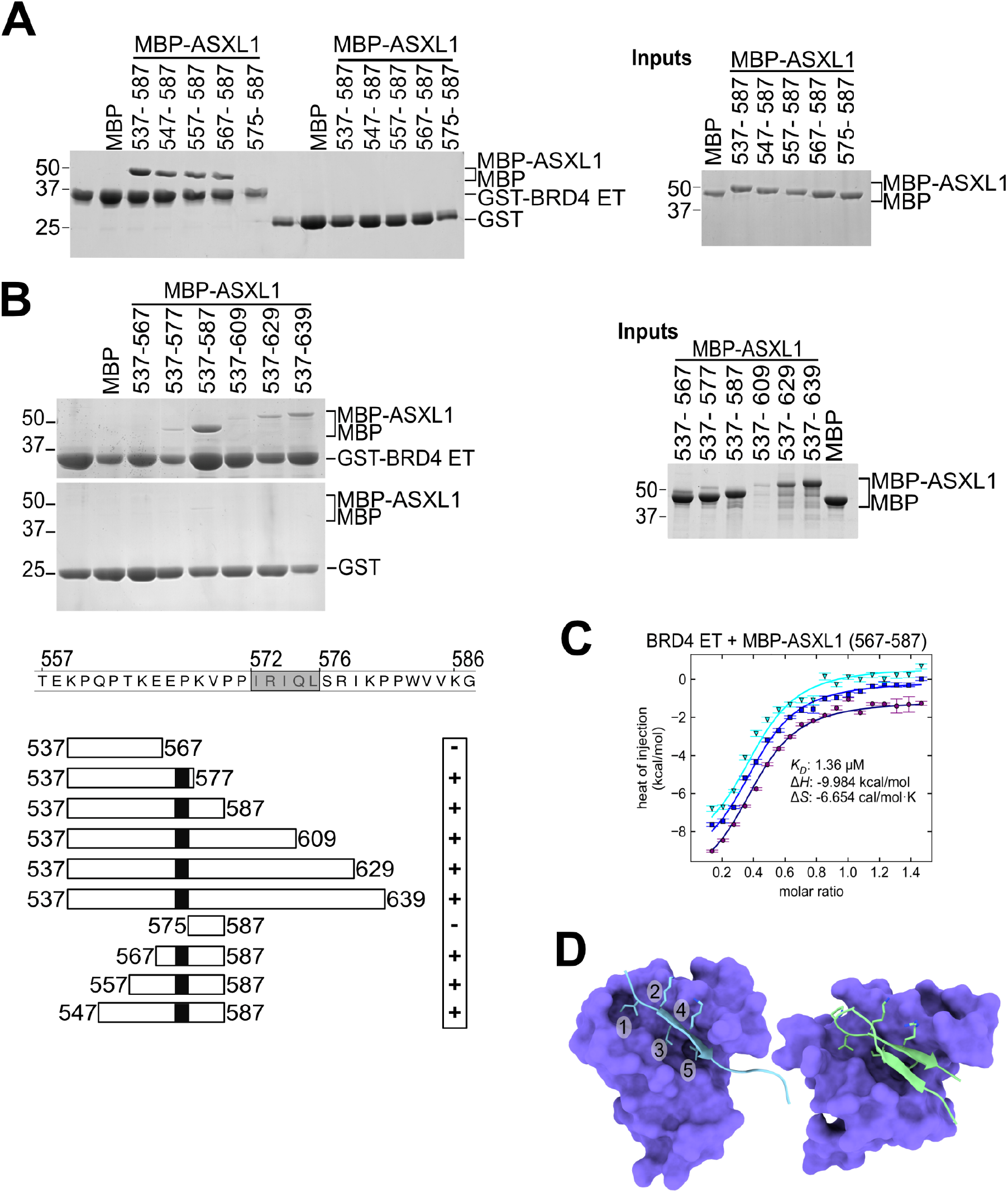
Refinement of the ASXL1 BRD4 binding site. Coomassie-stained gels showing pulldown of N-terminal (A) and C-terminal truncations (B) of His_6_MBP-ASXL1 by GST-BRD4-ET. Coomassie-stained gels to show relative expression of input His_6_MBP-ASXL1 constructs. (C) Isothermal titration calorimetry of ASXL1 and BRD4-ET. Isothermal titration calorimetry (ITC) of shorter His_6_MBP-fused ASXL1^567-587^ injected into BRD4-ET domain reveals a high affinity interaction, with K_D_= 1.36 μM. Data points of triplicate titrations, following His_6_MBP titration subtraction. (D) BRD4-ET bound to MLV(2N3K) and NSD3 (2NCZ).

**Supplementary Figure S3.**
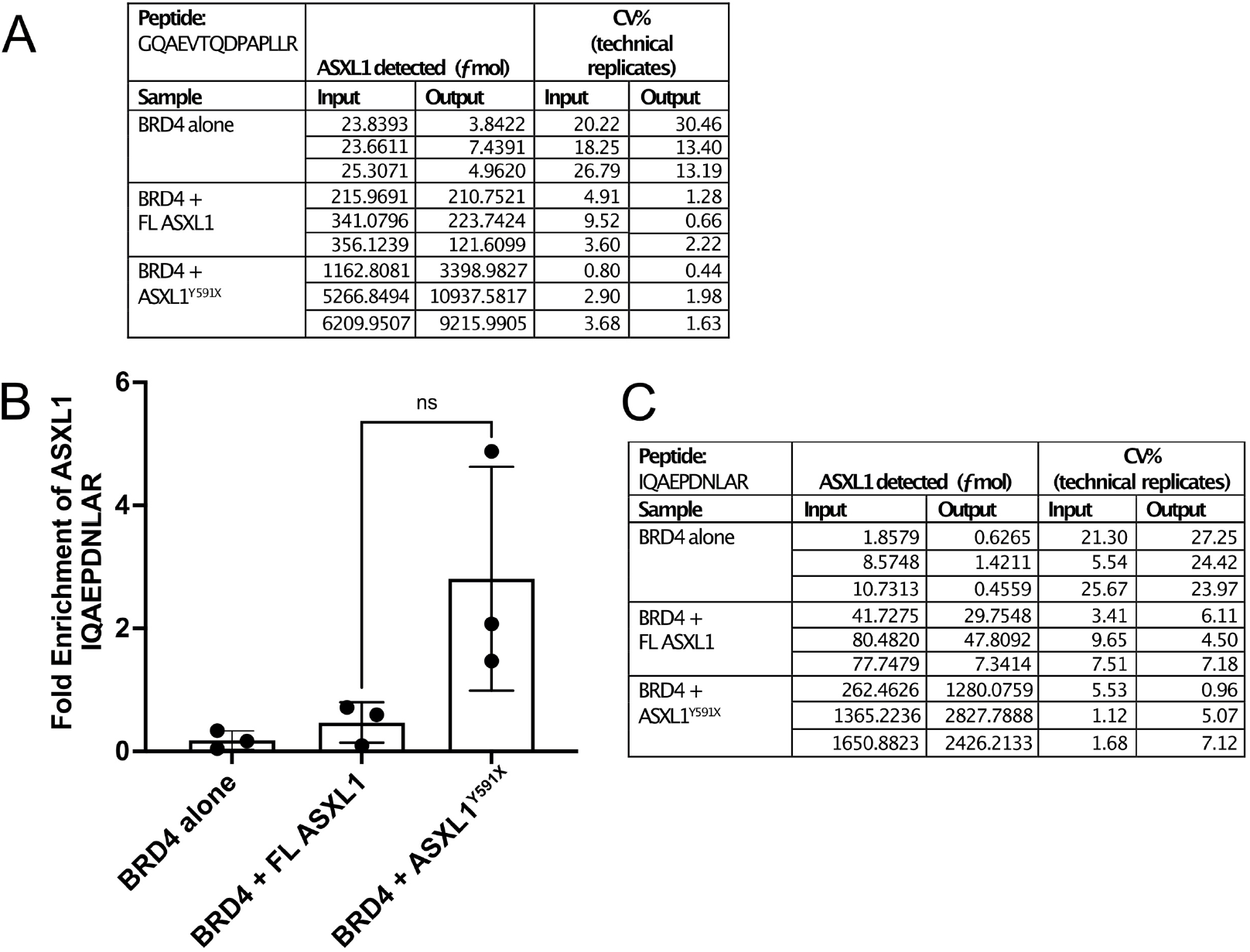
Quantification of ASXL1. Quantification of ASXL1 enrichment using parallel reaction-monitoring (PRM) mass spectrometry with isotopically-labelled peptide to determine fold enrichment of ASXL1 after one hour treatment with biotin and immunoprecipitation (A) Quantification using the second ASXL1 peptide (IQAEPDNLAR) (B) *f*mol of ASXL1 detected and CV % for IQAEPDNLAR (C) *f*mol of ASXL1 detected and CV % for GQAEVTQDPAPLLR, peptide analysis Fig4.

**Supplementary Figure S4.**
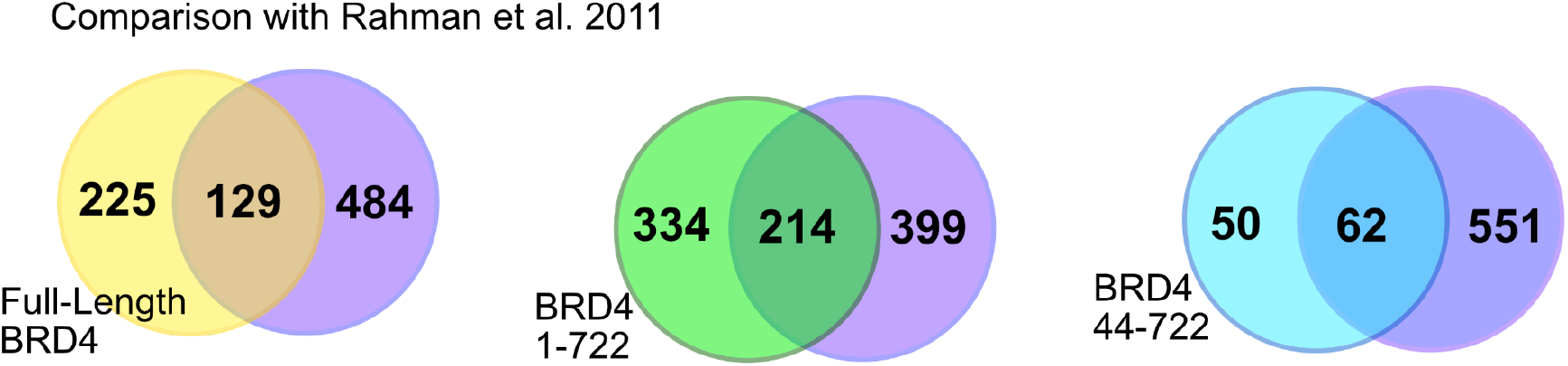
Comparison with Rahman et 2011. Overlap of proteins detected in our analysis (purple) with proteins detected after BRD4 IP by Rahman et al 2011. This paper used several different lengths of BRD4; Full-length (yellow), 1-722 (green) and 44-722 (blue). There is overlap with all the datasets and proteins detected from our analysis following enrichment with streptavidin of proteins biotinylated by TurboID-BRD4-ET.

**Supplementary Figure S5.**
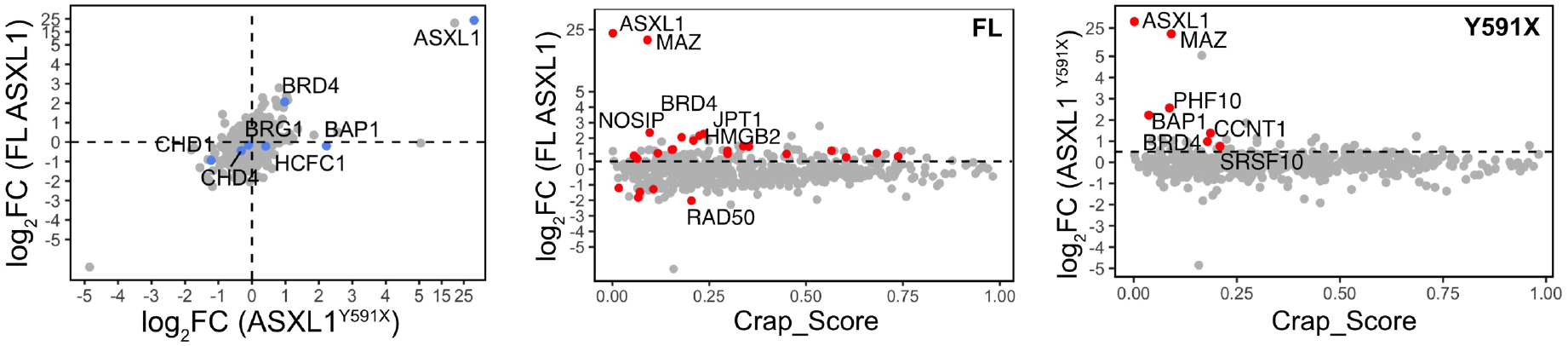
(A) Comparative plot of FL ASXL1 and ASXL1^Y591X^. The fold-change (ASXL1 FL/BRD4 alone) is plotted against the fold-change (ASXL1 ^Y591X^/BRD4 alone) to determine any difference in profile. Proteins which are known BRD4-ET interactors and PR-DUB proteins are highlighted in blue. The log2FC of each FL ASXL1 (B) and ASXL1^Y591X^ (C) (against BRD4-ET alone) was plotted against the crap_score. Proteins with significant p-values are highlighted (red). Proteins of interest are those which have a low crap_score but are significantly changed (highlighted in red)

## REFERENCES

Abdel-Wahab O, Adli M, LaFave LM, Gao J, Hricik T, Shih AH, Pandey S, Patel JP, Chung YR, Koche R, et al (2012) ASXL1 mutations promote myeloid transformation through loss of PRC2-mediated gene repression. Cancer Cell 22: 180–193

Asada S, Fujino T, Goyama S & Kitamura T (2019) The role of ASXL1 in hematopoiesis and myeloid malignancies. Cell Mol Life Sci 76: 2511–2523

Asada S, Goyama S, Inoue D, Shikata S, Takeda R, Fukushima T, Yonezawa T, Fujino T, Hayashi Y, Kawabata KC, et al (2018) Mutant ASXL1 cooperates with BAP1 to promote myeloid leukaemogenesis. Nat Commun 9: 2733

Bainbridge MN, Hu H, Muzny DM, Musante L, Lupski JR, Graham BH, Chen W, Gripp KW, Jenny K, Wienker TF, et al (2013) De novo truncating mutations in ASXL3 are associated with a novel clinical phenotype with similarities to Bohring-Opitz syndrome. Genome Med 5: 11

Balasubramani A, Larjo A, Bassein JA, Chang X, Hastie RB, Togher SM, Lähdesmäki H & Rao A (2015) Cancer-associated ASXL1 mutations may act as gain-of-function mutations of the ASXL1-BAP1 complex. Nat Commun 6: 7307

Bossone SA, Asselin C, Patel AJ & Marcu KB (1992) MAZ, a zinc finger protein, binds to c-MYC and C2 gene sequences regulating transcriptional initiation and termination. Proc Natl Acad Sci U S A 89: 7452–7456

Bott M, Brevet M, Taylor BS, Shimizu S, Ito T, Wang L, Creaney J, Lake RA, Zakowski MF, Reva B, et al (2011) The nuclear deubiquitinase BAP1 is commonly inactivated by somatic mutations and 3p21.1 losses in malignant pleural mesothelioma. Nat Genet 43: 668–672

Branon TC, Bosch JA, Sanchez AD, Udeshi ND, Svinkina T, Carr SA, Feldman JL, Perrimon N & Ting AY (2018) Efficient proximity labeling in living cells and organisms with TurboID. Nat Biotechnol 36: 880–887

Carbone M, Yang H, Pass HI, Krausz T, Testa JR & Gaudino G (2013) BAP1 and cancer. Nat Rev Cancer 13: 153–159

Chittock EC, Latwiel S, Miller TCR & Müller CW (2017) Molecular architecture of polycomb repressive complexes. Biochem Soc Trans 45: 193–205

Conrad RJ, Fozouni P, Thomas S, Sy H, Zhang Q, Zhou M-M & Ott M (2017) The short isoform of BRD4 promotes HIV-1 latency by engaging repressive SWI/SNF chromatin-remodeling complexes. Mol Cell 67: 1001–1012.e6

Crowe BL, Larue RC, Yuan C, Hess S, Kvaratskhelia M & Foster MP (2016) Structure of the Brd4 ET domain bound to a C-terminal motif from γ-retroviral integrases reveals a conserved mechanism of interaction. Proc Natl Acad Sci U S A 113: 2086–2091

Daou S, Barbour H, Ahmed O, Masclef L, Baril C, Sen Nkwe N, Tchelougou D, Uriarte M, Bonneil E, Ceccarelli D, et al (2018) Monoubiquitination of ASXLs controls the deubiquitinase activity of the tumor suppressor BAP1. Nat Commun 9: 4385

Dawson MA, Prinjha RK, Dittmann A, Giotopoulos G, Bantscheff M, Chan W-I, Robson SC, Chung C-W, Hopf C, Savitski MM, et al (2011) Inhibition of BET recruitment to chromatin as an effective treatment for MLL-fusion leukaemia. Nature 478: 529–533

De I, Chittock EC, Grötsch H, Miller TCR, McCarthy AA & Müller CW (2018) Structural Basis for the Activation of the Deubiquitinase Calypso by the Polycomb Protein ASX. Structure

Devaiah BN, Case-Borden C, Gegonne A, Hsu CH, Chen Q, Meerzaman D, Dey A, Ozato K & Singer DS (2016) BRD4 is a histone acetyltransferase that evicts nucleosomes from chromatin. Nat Struct Mol Biol 23: 540–548

Dey A, Nishiyama A, Karpova T, McNally J & Ozato K (2009) Brd4 marks select genes on mitotic chromatin and directs postmitotic transcription. Mol Biol Cell 20: 4899–4909

Dey A, Seshasayee D, Noubade R, French DM, Liu J, Chaurushiya MS, Kirkpatrick DS, Pham VC, Lill JR, Bakalarski CE, et al (2012) Loss of the tumor suppressor BAP1 causes myeloid transformation. Science 337: 1541–1546

Duan Y, Guan Y, Qin W, Zhai X, Yu B & Liu H (2018) Targeting Brd4 for cancer therapy: inhibitors and degraders. Medchemcomm 9: 1779–1802

Foglizzo M, Middleton AJ, Burgess AE, Crowther JM, Dobson RCJ, Murphy JM, Day CL & Mace PD (2018) A bidentate Polycomb Repressive-Deubiquitinase complex is required for efficient activity on nucleosomes. Nat Commun 9: 3932

Gao J, Aksoy BA, Dogrusoz U, Dresdner G, Gross B, Sumer SO, Sun Y, Jacobsen A, Sinha R, Larsson E, et al (2013) Integrative analysis of complex cancer genomics and clinical profiles using the cBioPortal. Sci Signal 6: l1

Gelsi-Boyer V, Brecqueville M, Devillier R, Murati A, Mozziconacci M-J & Birnbaum D (2012) Mutations in ASXL1 are associated with poor prognosis across the spectrum of malignant myeloid diseases. J Hematol Oncol 5: 12

Hoischen A, van Bon BWM, Rodríguez-Santiago B, Gilissen C, Vissers LELM, de Vries P, Janssen I, van Lier B, Hastings R, Smithson SF, et al (2011) De novo nonsense mutations in ASXL1 cause Bohring-Opitz syndrome. Nat Genet 43: 729–731

Inoue D, Kitaura J, Togami K, Nishimura K, Enomoto Y, Uchida T, Kagiyama Y, Kawabata KC, Nakahara F, Izawa K, et al (2013) Myelodysplastic syndromes are induced by histone methylation–altering ASXL1 mutations. J Clin Invest 123: 4627–4640

Inoue D, Matsumoto M, Nagase R, Saika M, Fujino T, Nakayama KI & Kitamura T (2016) Truncation mutants of ASXL1 observed in myeloid malignancies are expressed at detectable protein levels. Exp Hematol 44: 172–6.e1

Jordan-Sciutto KL, Dragich JM, Caltagarone J, Hall DJ & Bowser R (2000) Fetal Alz-50 clone 1 (FAC1) protein interacts with the Myc-associated zinc finger protein (ZF87/MAZ) and alters its transcriptional activity. Biochemistry 39: 3206–3215

Käll L, Canterbury JD, Weston J, Noble WS & MacCoss MJ (2007) Semi-supervised learning for peptide identification from shotgun proteomics datasets. Nat Methods 4: 923–925

Lambert J-P, Picaud S, Fujisawa T, Hou H, Savitsky P, Uusküla-Reimand L, Gupta GD, Abdouni H, Lin Z-Y, Tucholska M, et al (2019) Interactome Rewiring Following Pharmacological Targeting of BET Bromodomains. Mol Cell 73: 621–638.e17

Laugesen A, Højfeldt JW & Helin K (2019) Molecular Mechanisms Directing PRC2 Recruitment and H3K27 Methylation. Mol Cell 74: 8–18

Mann M, Roberts DS, Zhu Y, Li Y, Zhou J, Ge Y & Brasier AR (2021) Discovery of RSV-induced BRD4 protein interactions using native immunoprecipitation and Parallel Accumulation-Serial Fragmentation (PASEF) mass spectrometry. Viruses 13: 454

Mellacheruvu D, Wright Z, Couzens AL, Lambert J-P, St-Denis NA, Li T, Miteva YV, Hauri S, Sardiu ME, Low TY, et al (2013) The CRAPome: a contaminant repository for affinity purification-mass spectrometry data. Nat Methods 10: 730–736

Micol J-B, Pastore A, Inoue D, Duployez N, Kim E, Lee SC-W, Durham BH, Chung YR, Cho H, Zhang X-J, et al (2017) ASXL2 is essential for haematopoiesis and acts as a haploinsufficient tumour suppressor in leukemia. Nat Commun 8: 15429

Mochizuki K, Nishiyama A, Jang MK, Dey A, Ghosh A, Tamura T, Natsume H, Yao H & Ozato K (2008) The bromodomain protein Brd4 stimulates G1 gene transcription and promotes progression to S phase. J Biol Chem 283: 9040–9048

Nagase R, Inoue D, Pastore A, Fujino T, Hou H-A, Yamasaki N, Goyama S, Saika M, Kanai A, Sera Y, et al (2018) Expression of mutant Asxl1 perturbs hematopoiesis and promotes susceptibility to leukemic transformation. J Exp Med 215: 1729–1747

Rahman S, Sowa ME, Ottinger M, Smith JA, Shi Y, Harper JW & Howley PM (2011) The Brd4 extraterminal domain confers transcription activation independent of pTEFb by recruiting multiple proteins, including NSD3. Mol Cell Biol 31: 2641–2652

Reddington CJ, Fellner M, Burgess AE & Mace PD (2020) Molecular Regulation of the Polycomb Repressive-Deubiquitinase. Int J Mol Sci 21

Sahtoe DD, van Dijk WJ, Ekkebus R, Ovaa H & Sixma TK (2016) BAP1/ASXL1 recruitment and activation for H2A deubiquitination. Nat Commun 7: 10292

Scheuermann JC, de Ayala Alonso AG, Oktaba K, Ly-Hartig N, McGinty RK, Fraterman S, Wilm M, Muir TW & Müller J (2010) Histone H2A deubiquitinase activity of the Polycomb repressive complex PR-DUB. Nature 465: 243–247

Shashi V, Pena LDM, Kim K, Burton B, Hempel M, Schoch K, Walkiewicz M, McLaughlin HM, Cho M, Stong N, et al (2016) De Novo truncating variants in ASXL2 are associated with a unique and recognizable clinical phenotype. Am J Hum Genet 99: 991–999

Shen C, Ipsaro JJ, Shi J, Milazzo JP, Wang E, Roe J-S, Suzuki Y, Pappin DJ, Joshua-Tor L & Vakoc CR (2015) NSD3-Short Is an Adaptor Protein that Couples BRD4 to the CHD8 Chromatin Remodeler. Mol Cell 60: 847–859

Shukla V, Rao M, Zhang H, Beers J, Wangsa D, Wangsa D, Buishand FO, Wang Y, Yu Z, Stevenson HS, et al (2017) ASXL3 is a novel pluripotency factor in human respiratory epithelial cells and a potential therapeutic target in small cell lung cancer. Cancer Res 77: 6267–6281

Silva JC, Gorenstein MV, Li G-Z, Vissers JPC & Geromanos SJ (2006) Absolute quantification of proteins by LCMSE: a virtue of parallel MS acquisition. Mol Cell Proteomics 5: 144–156

Srivastava A, Ritesh KC, Tsan Y-C, Liao R, Su F, Cao X, Hannibal MC, Keegan CE, Chinnaiyan AM, Martin DM, et al (2016) De novo dominant ASXL3 mutations alter H2A deubiquitination and transcription in Bainbridge-Ropers syndrome. Hum Mol Genet 25: 597–608

Szczepanski AP, Zhao Z, Sosnowski T, Goo YA, Bartom ET & Wang L (2020) ASXL3 bridges BRD4 to BAP1 complex and governs enhancer activity in small cell lung cancer. Genome Med 12: 63

Testa JR, Cheung M, Pei J, Below JE, Tan Y, Sementino E, Cox NJ, Dogan AU, Pass HI, Trusa S, et al (2011) Germline BAP1 mutations predispose to malignant mesothelioma. Nat Genet 43: 1022–1025

Wai DCC, Szyszka TN, Campbell AE, Kwong C, Wilkinson-White LE, Silva APG, Low JKK, Kwan AH, Gamsjaeger R, Chalmers JD, et al (2018) The BRD3 ET domain recognizes a short peptide motif through a mechanism that is conserved across chromatin remodelers and transcriptional regulators. J Biol Chem 293: 7160–7175

Wang L, Birch NW, Zhao Z, Nestler CM, Kazmer A, Shilati A, Blake A, Ozark PA, Rendleman EJ, Zha D, et al (2021) Epigenetic targeted therapy of stabilized BAP1 in ASXL1 gain-of-function mutated leukemia. Nature Cancer 2: 515–526

Wiesner T, Obenauf AC, Murali R, Fried I, Griewank KG, Ulz P, Windpassinger C, Wackernagel W, Loy S, Wolf I, et al (2011) Germline mutations in BAP1 predispose to melanocytic tumors. Nat Genet 43: 1018–1021

Wu T, Kamikawa YF & Donohoe ME (2018) Brd4’s bromodomains mediate histone H3 acetylation and chromatin remodeling in pluripotent cells through P300 and Brg1. Cell Rep 25: 1756–1771

Yang H, Kurtenbach S, Guo Y, Lohse I, Durante MA, Li J, Li Z, Al-Ali H, Li L, Chen Z, et al (2018) Gain of function of ASXL1 truncating protein in the pathogenesis of myeloid malignancies. Blood 131: 328–341

Yang Z, He N & Zhou Q (2008) Brd4 recruits P-TEFb to chromosomes at late mitosis to promote G1 gene expression and cell cycle progression. Mol Cell Biol 28: 967–976

Zhang P, He F, Bai J, Yamamoto S, Chen S, Zhang L, Sheng M, Zhang L, Guo Y, Man N, et al (2018) Chromatin regulator Asxl1 loss and Nf1 haploinsufficiency cooperate to accelerate myeloid malignancy. J Clin Invest 128: 5383–5398

Zhang Q, Zeng L, Shen C, Ju Y, Konuma T, Zhao C, Vakoc CR & Zhou M-M (2016) Structural Mechanism of Transcriptional Regulator NSD3 Recognition by the ET Domain of BRD4. Structure 24: 1201–1208

